# Efficient targeting of NY-ESO-1 tumor antigen to human cDC1s by lymphotactin results in cross-presentation and antigen-specific T cell expansion

**DOI:** 10.1101/2021.11.30.470538

**Authors:** Camille M. Le Gall, Anna Cammarata, Lukas de Haas, Iván Ramos-Tomillero, Jorge Cuenca-Escalona, Zacharias Wijfjes, Anouk M.D. Becker, Yusuf Dölen, I. Jolanda M. de Vries, Carl G. Figdor, Georgina Flórez-Grau, M. Verdoes

**Affiliations:** Department of Tumor Immunology, Radboud Institute for Molecular Life Sciences, Radboudumc, Geert Grooteplein Zuid 28, 6525 GA, Nijmegen, The Netherlands; Oncode Institute, Geert Grooteplein Zuid 28, 6525 GA, Nijmegen, The Netherlands; Institute for Chemical Immunology, Geert Grooteplein Zuid 28, 6525 GA, Nijmegen, Netherlands

## Abstract

Type 1 conventional dendritic cells (cDC1s) are characterized by their ability to induce potent CD8^+^ T cell responses. In efforts to generate novel vaccination strategies, notably against cancer, human cDC1s emerge as an ideal target to deliver antigens. cDC1s uniquely express XCR1, a seven transmembrane G protein-coupled receptor (GPCR). Due to its restricted expression and endocytic nature, XCR1 represents an attractive receptor to mediate antigen-delivery to human cDC1s. To explore tumor antigen delivery to human cDC1s, we used an engineered version of XCR1-binding lymphotactin (XCL1), XCL1(CC3). Site-specific sortase-mediated transpeptidation was performed to conjugate XCL1(CC3) to an analog of the HLA-A*02:01 epitope of the cancer testis antigen New York Esophageal Squamous Cell Carcinoma-1 (NY-ESO-1). While poor epitope solubility prevented isolation of stable XCL1-antigen conjugates, incorporation of a single polyethylene glycol (PEG) chain upstream of the epitope-containing peptide enabled generation of soluble XCL1(CC3)-antigen fusion constructs. Binding and chemotactic characteristics of the XCL1-antigen conjugate, as well as its ability to induce antigen-specific CD8^+^ T cell activation by cDC1s, was assessed. PEGylated XCL1(CC3)-antigen conjugates retained binding to XCR1, and induced cDC1 chemoattraction *in vitro*. The model epitope was efficiently cross-presented by human cDC1s to activate NY-ESO-1-specific CD8^+^ T cells. Importantly, vaccine activity was increased by targeting XCR1 at the surface of cDC1s. Our results present a novel strategy for the generation of targeted vaccines fused to insoluble antigens. Moreover, our data emphasize the potential of targeting XCR1 at the surface of primary human cDC1s to induce potent CD8^+^ T cell responses.

## INTRODUCTION

Conventional DCs type 1 (cDC1s) are the rarest subset of dendritic cells (DCs), making up only 0.03% of human peripheral blood mononuclear cells (PBMCs). cDC1s are characterized by their expression of CD141 (BDCA-3), CLEC9A (DNGR-1) and X-C motif chemokine receptor 1 (XCR1)^1, 2^, and additionally express high levels of Toll-like receptor (TLR) 3 and TLR7. The scarcity of these cells in peripheral blood renders the study of human cDC1s cumbersome, thus most functional studies have been performed on their murine cDC1s counterpart. *Batf3*-deficient mice selectively lacking cDC1s are unable to mount an effective cytotoxic immune response against viruses and tumors^3^. cDC1s have been shown *in vitro* and *in vivo* to excel at cross-presentation of extracellular^4^ and dead-cell associated^5^ antigens to CD8^+^ T cells. Human CD141^+^CLEC9A^+^XCR1^+^ have been shown to similarly excel at cross-presentation^6, 7^. This cross-presenting capacity makes human cDC1s an optimal cell population for eliciting cytotoxic immune responses against tumors.

cDC1s uniquely express XCR1, a chemokine receptor allowing cells to specifically migrate towards lymphotactin, commonly referred to as X-C motif chemokine ligand 1 (XCL1). XCL1 is a 12 kDa chemokine mainly secreted by activated cytotoxic CD8^+^ T and NK cells. The XCR1/XCL1 axis is a major regulator of cytotoxic immune responses. Mice lacking either XCR1 or XCL1 have deficient cytotoxic T cell responses^8^, but interestingly also lack the ability to generate regulatory T cells^9^. XCL1 is thus able to modulate the spatial location and function of both T cells and DCs. Binding of XCL1 to the orthosteric site of XCR1 triggers Ca^2+^ efflux from the endoplasmic reticulum (ER), leading to cytoskeleton remodeling and cell migration^10^, followed by XCR1 desensitization and internalization to early endosomes^11^. It is hypothesized that by following a XCL1 chemotactic gradient, XCR1^+^ cDC1s can migrate towards the site of inflammation, where they take up antigens. Downregulation of XCR1, and activation-induced expression of CCR7, enables subsequent migration to the lymph node^12^, where cDC1s are able to prime T cells. Due to the restricted expression of XCR1 on cDC1s, and its endocytic nature, XCR1 represents a highly attractive target for the delivery of tumor antigens *in vivo*, and to induce CD8^+^ T cell responses^13–15^. We chose to use its ligand, XCL1, as a tumor antigen delivery moiety^14, 16, 17^.

XCL1 is the only member of its family in mouse. In humans, XCL2 (NC_000001.11 (168540768..168543997, complement)) is a paralog chemokine present in the same locus as XCL1 (NC_000001.11 (168574128..168582069)). XCL1 and XCL2 have the particularity to present only one disulfide bond. This particularity allows them to adopt two conformations under normal physiological conditions: a monomeric α-β XCR1-binding fold, and an all-β dimeric glycosaminoglycan (GAG)-binding fold^18, 19^. Point mutations of two valine residues into cysteine residues (V21C/V59C) locks XCL1 into its XCR1 agonist α-β fold^19, 20^. We generated an engineered XCR1 ligand, based on the V21C / V59C fold (XCL1(CC3)), further modified by fusing it C-terminally to a LPETGG sortag motif, allowing for site-specific chemoenzymatic modification to virtually any payload^21^.

To build and evaluate the activity of a cDC1-targeting vaccine, we set out to conjugate XCL1(CC3) to an epitope of the well-characterized tumor antigen New York Esophageal Squamous Cell Carcinoma-1 (NY-ESO-1). This cancer testis tumor-associated antigen (TAA) is aberrantly expressed by a large proportion of patients in multiple malignancies, including multiple myeloma (60%)^22^, neuroblastoma (82%)^23^, and melanoma (45%)^24^. NY-ESO-1-derived peptide (157-165) (^157^SLLMWITQ^165^C, in short S7C) has been identified as an immunodominant epitope giving rise to antigen-specific CD8^+^ T cell responses in HLA-A*02:01 patients^25^. S7C is notoriously highly hydrophobic (hydrophobicity = 35.77, GRAVY = 1.18), and dimer formation by P9 cysteine-pairing causes problems in preparation and formulation of S7C-based vaccines^26^. Despite their unfavorable biochemical characteristics, NY-ESO-1-derived epitopes are highly promising for off-the-shelf cancer vaccines, due to NY-ESO-1 high prevalence. To avoid cysteine pairing, we chose to use an analog of S7C as model antigen (^157^SLLMWITQ(^165^Abu) (S7Abu), Abu = L-2-aminobutyric acid), which was shown to bind HLA-A*02:01, and to be recognized by S7C-specific patient-derived CD8^+^ T cells^26^. We aimed at functionalizing XCL1 with the S7Abu epitope, and evaluating the potential of such a targeting approach to activate antigen-specific CD8^+^ T cells via human XCR1^+^ cDC1s.

## RESULTS

### XCL1(CC3) can be site-specifically labeled without disrupting binding to XCR1

In order to site-specifically modify XCL1(CC3), we genetically fused it to a *N*-terminal His-tagged SUMO solubility tag, a LPETGG sortag motif, and a FLAG tag (figure 1A). After production and refolding, we isolated XCL1(CC3)-LPETGG-FLAG by nickel affinity purification (figure 1B). This typically yielded ∼5-10 mg XCL1(CC3) per liter of starting culture, with a refolding efficacy over 75%, as analyzed by reverse phase HPLC (figure S1A). To confirm that XCL1(CC3) was able to retain its binding specificity following site-directed modification, we subjected XCL1(CC3)-LPETGG-FLAG to site-specific conjugation to a small fluorophore-equipped peptide [H-GGGCK(FITC)-NH_2_] (referred to as FITC). We were able to isolate XCL1(CC3)-FITC with a yield of ∼45% (figure S1B). To confirm binding capacity and specificity, we incubated a mixture of freshly isolated cDC2s (CD1c^+^CD141^-^XCR1^-^) and cDC1s (CD14^+^XCR1^+^), with either a commercial anti-XCR1 antibody or XCL1(CC3)-FITC. As analyzed by flow cytometry (figures 1C and S1C), XCL1(CC3)-FITC specifically identified a subpopulation of cDC1s (∼60% of cDC1s), similarly to the commercial antibody. Moreover, XCR1 expression on primary cDC1s could be visualized by confocal microscopy using XCL1(CC3)-Cy5.5 (figure 1D), which was generated similarly. Hence, introduction of a LPETGG motif at the C-terminus of XCL1(CC3) did not impair folding, and the engineered chemokine could be modified site-specifically with fluorescent peptides, while retaining its binding capacity.

**Figure 1:**
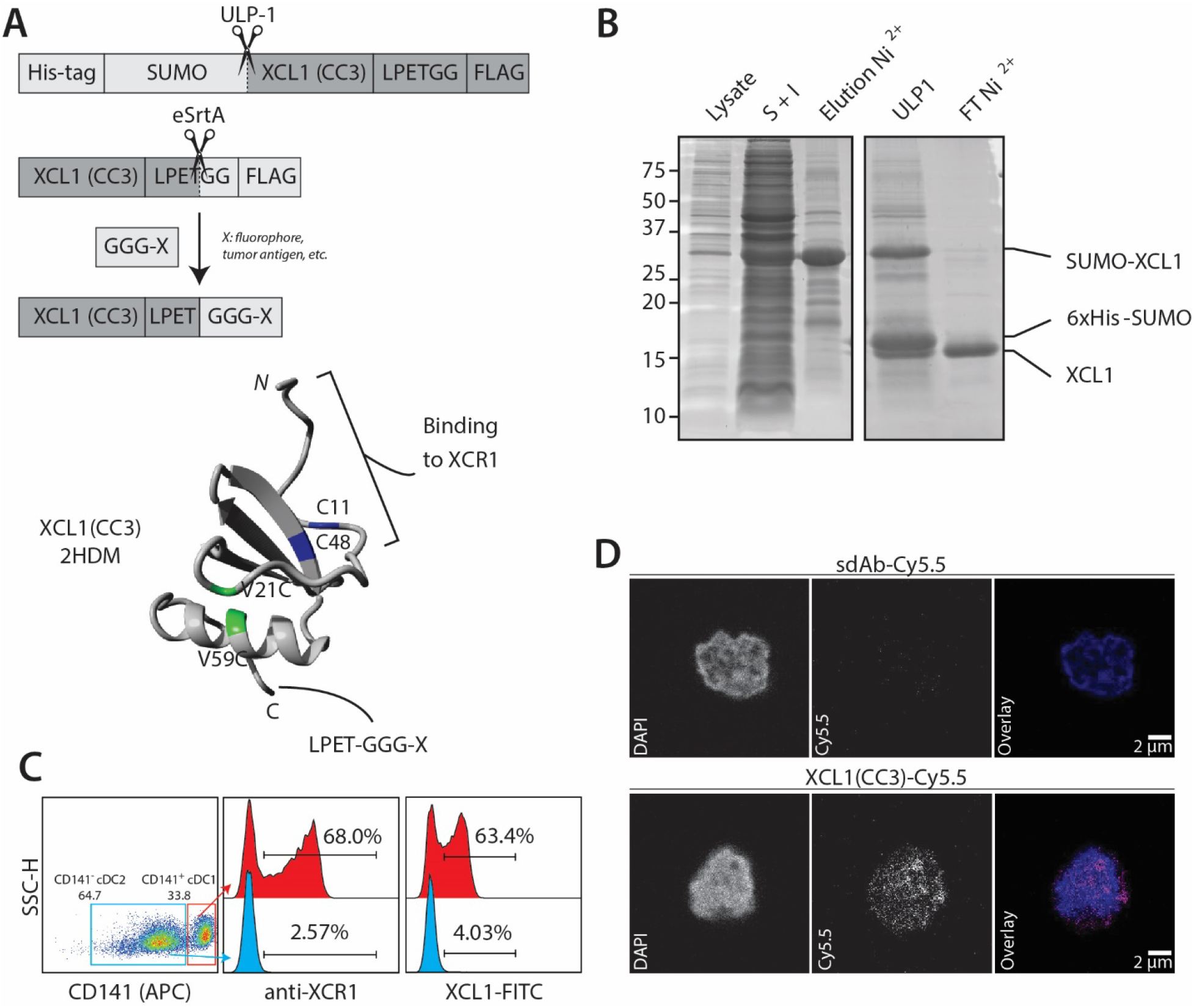
XCL1(CC3)–LPETG retains binding specificity to XCR1 after site-specific labeling with a small fluorophore. **A.** Representation of 6×His-SUMO-XCL1-LPETGG-FLAG produced in BL21(DE3), site-specific labelling via sortase-mediated transpeptidation, and structure of XCL1(CC3) after purification and refolding. V21C and V59C point mutations stabilizing the α-β chemokine fold are highlighted in green, and two cysteine residues C11 and C48 present in the native XCL1 are highlighted in blue. Adapted from 2HDM structure and modeled in YASARA. **B.** Production of XCL1(CC3) in BL21(DE3). ∼30 kDa 6×His-SUMO-XCL1(CC3)-LPETGG-FLAG is present in bacterial lysate (lane 1) and enriched by pooling the soluble and insoluble fractions (S+I, lane 2). 6×His-SUMO-XCL1 is eluted (lane 3) after nickel affinity purification, and solubility tag digested (ULP1, lane 4). XCL1(CC3) is isolated after a second nickel purification (FT Ni^2+^, lane 5) with high purity. **C.** Commercial anti-XCR1-PE and XCL1(CC3)-FITC identify a comparable subpopulation of ∼60% CD141^+^ cDC1s without staining CD141^-^ cDC2s, confirming that XCL1(CC3)-FITC specifically binds to XCR1. **D.** Confocal microscopy performed on cDC1s with XCL1(CC3)-Cy5.5 or sdAb-Cy5.5 shows specificity of XCL1(CC3) for XCR1.

### Poor solubility of tumor antigen prevents conjugation to XCL1

After confirming that XCL1(CC3)-LPETGG-FLAG could be modified while retaining XCR1 binding capacity, we set out to generate a construct able to deliver tumor antigens to cDC1s. To this end, we synthesized a peptide containing a triglycine motif, an azido-lysine as clickable handle, a FR-dipeptide motif to promote cross-presentation by biasing proteasomal cleavage^27^, followed by S7Abu, ([GGGK(N_3_)FRSLLMWITQ(Abu)], referred to as K(N_3_)-S7Abu). Upon site-specific labeling of XCL1(CC3)-LPETGG-FLAG, we detected formation of XCL1(CC3)-K(N_3_)-S7Abu with minimal hydrolysis product (XCL1-H) (figure S1D). However, the product could not be purified (figure S2A), suggesting that XCL1-K(N_3_)-S7Abu was not stable in solution. Further analysis (cf. methods) indicated that the product was entirely present in aggregates formed during the reaction (figure S2B). Sortase-mediated transpeptidation exchanged a solubilizing His-tag for an extremely hydrophobic S7Abu epitope, likely causing this aggregation.

### PEGylation of tumor antigen allows conjugation of S7Abu to XCL1(CC3)

As peptide conjugation appeared to induce aggregation of XCL1(CC3)-K(N_3_)-S7Abu, we decided to use the azido-lysine to introduce a 5 kDa polyethylene glycol (PEG5k) chain via copper free strain-promoted azide–alkyne cycloaddition (SPAAC) (figure 2A). We performed a SPAAC reaction of [K(N_3_)-S7Abu] with a three-fold excess of DBCO-PEG5k. Reaction completion was confirmed by HPLC (figure S2C) and Matrix-assisted laser desorption/ionization-Time-of-Flight (MALDI-TOF) (figure S2D). Excess DBCO-PEG5k was capped by addition of NaN_3_, followed by site-specific conjugation of [GGGK(PEG5k)FR-S7Abu] (K(PEG)-S7Abu) to XCL1. We observed formation of XCL1(CC3)-K(PEG)-S7Abu, which we could purify by cation exchange chromatography with an isolated yield above 40%, and over 75% purity as seen by SDS-PAGE (figure 2B). A competitive flow cytometry-based binding assay performed with freshly isolated cDC1s by flow cytometry revealed that presence of XCL1(CC3)-K(PEG)-S7Abu was able to prevent binding of a XCR1 commercial antibody (figure 2C). Thus, modification and PEGylation of the chemokine did not impair its binding capacity. Importantly, XCL1(CC3)-K(PEG)-S7Abu also had similar chemotactic activity compared to commercial XCL1, and unmodified XCL1(CC3) (figure 2D), demonstrating that modification of the chemokine did not affect its functionality. In lieu of an isotype control, we generated a non-specific vaccine by modifying an in-house-generated single domain antibody (sdAb) directed against BDCA-2 (CD303), a plasmacytoid DC marker, and could isolate sdAb-K(PEG)-S7Abu using a similar method, in comparable yields (figure S2E).

**Figure 2:**
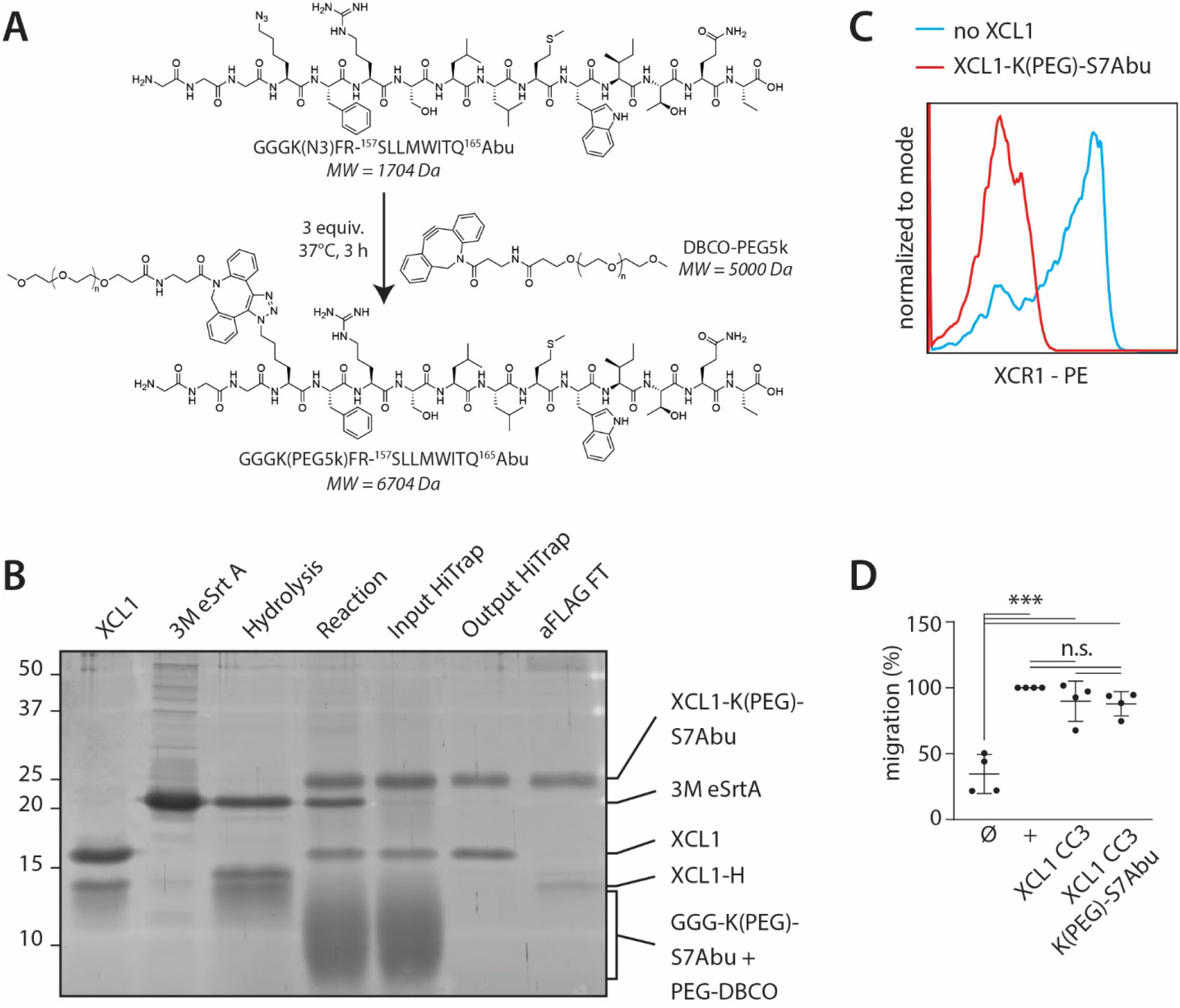
PEGylation of GGG-K(N_3_)-S7Abu enables its conjugation to XCL1(CC3). **A.** Site-specific PEGylation of GGG-K(N_3_)-S7Abu with DBCO-PEG5k. **B.** Site-specific labeling of XCL1(CC3) (lane 1) with GGG-K(PEG)-S7Abu allows product formation (lane 4). Excess of nucleophile and PEG5k is removed after cation exchange (lane 6), and unreacted XCL1(CC3) removed by incubation with anti-FLAG beads, allowing isolation of pure XCL1(CC3)-K(PEG)-S7Abu (lane 7). 3M eSrtA (lane 2) activity is confirmed by hydrolysis of XCL1(CC3) in absence of peptide (lane 3). Densitometry was performed to calculate the concentration of XCL1(CC3)-K(PEG)-S7Abu used in cell experiments. **C.** Incubation of XCR1-expressing cDC1s with anti-XCR1 in presence or absence of XCL1(CC3)-K(PEG)-S7Abu shows that the PEGylated vaccine retains binding to XCR1. **D.** cDC1 migration towards media (Ø), 10 ng.mL^-1^ rhXCL1 (+, BioLegend), XCL1(CC3) and XCL1(CC3)-K(PEG)-S7Abu shows that modification of XCL1 does not impair its chemotactic activity. N=4 independent donors, normalized to rhXCL1. One-way ANOVA, * p<0.05, ** p<0.01, ***p<0.001.

### XCL1(CC3)-K(PEG)-S7Abu elicits potent CD8^+^ T cell responses following XCR1 mediated uptake by cDC1s

We analyzed the ability of primary human cDC1s exposed to XCL1(CC3)-K(PEG)-S7Abu to induce CD8^+^ T cell proliferation and activation. To analyze the influence of XCR1 targeting, we also treated cDC1s with an unspecific vehicle (sdAb-K(PEG)-S7Abu) (figure 3A). CD8^+^ T cells proliferated in response to presentation of their cognate antigen S7Abu on HLA-A*02:01 molecules in all conditions (figures 3B and S3A). cDC1s treated with 0.1 µM XCL1(CC3)-K(PEG)-S7Abu induced an increased CD8^+^ T cell proliferation compared to the control vehicle, as quantified by measuring CD8^+^ T cell division index (figure 3B). In addition, CD8^+^ T cells activated by cDC1s treated with 0.1 µM XCL1(CC3)-K(PEG)-S7Abu displayed an increased expression of IL2Rα (CD25), downregulated Interleukin-7 (IL-7) receptor (IL-7R) and T cell immunoreceptor with Ig and ITIM domains (TIGIT) (figures 3C and S3B), and secreted increased levels of IL-2 and interferon γ (IFNγ) (figure 3D). Similarly, increased PD-1 and CD137 levels were detected, but this did not reach statistical significance (figures 3C and S3B). cDC1s treated with higher doses (1 µM) of both XCR1-targeted and control constructs induced similar CD8^+^ T cell proliferation, and activation (figures 3B-D). These results showed that targeting of XCR1 increases tumor antigen uptake by cDC1s, resulting in an improved CD8^+^ T cell activation.

**Figure 3:**
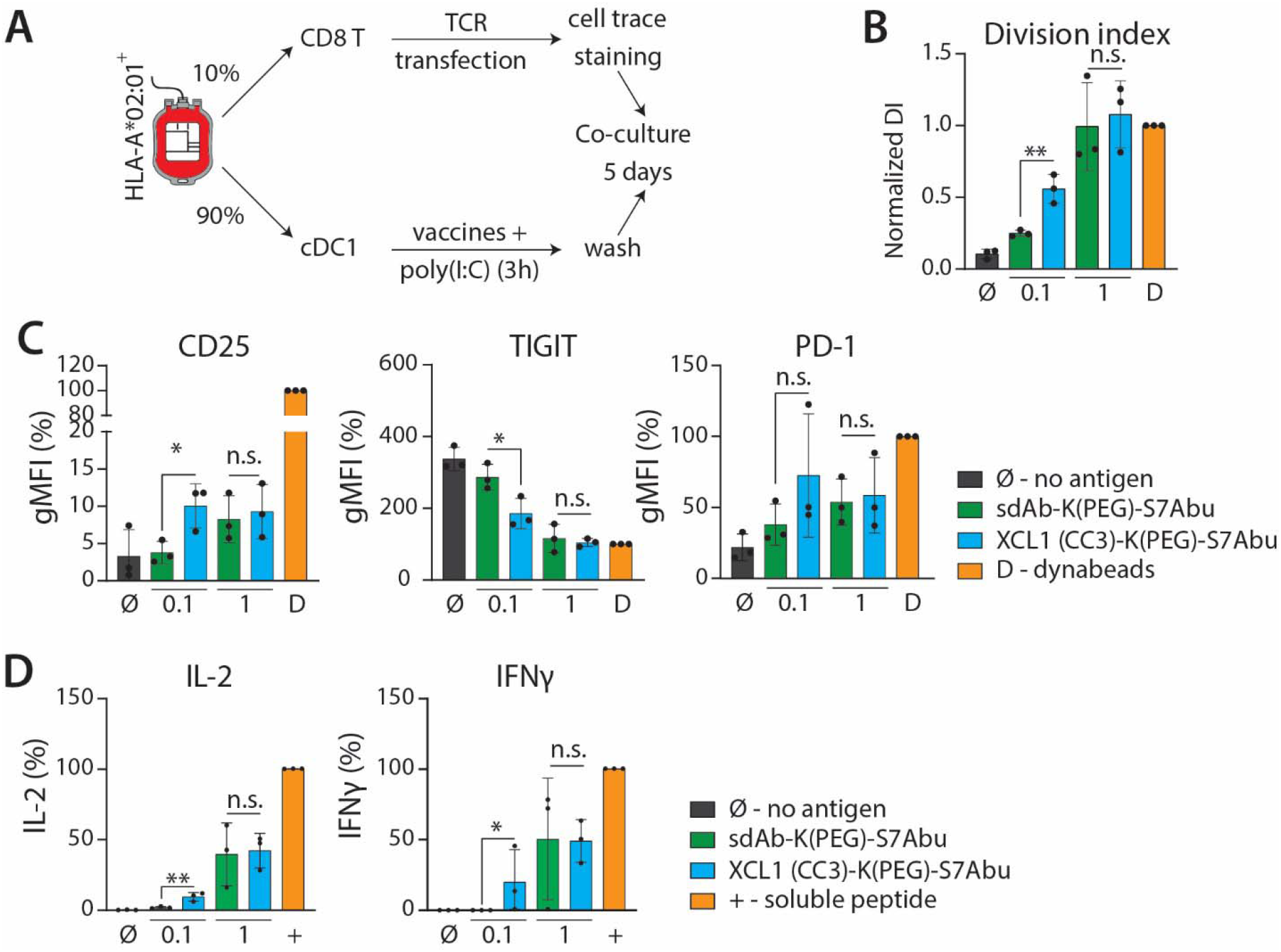
cDC1s treated with XCL1(CC3)-K(PEG)-S7Abu cross present S7Abu to CD8^+^ T cells following XCR1-mediated uptake. **A.** Experimental setup allowing for cDC1 isolation and CD8^+^ T cell transfection with S7Abu-specific TCR. **B.**, **C. and D.** Increased activation of CD8^+^ T cells through XCR1 targeting by cDC1s treated with XCL1(CC3)-K(PEG)-S7Abu compared to vehicle control, as measured by CD8^+^ T cell division index (B), expression of activation markers (CD25, PD-1) and downregulation of TIGIT (C), and IL-2 and IFNγ secretion in the culture media (D). N=3 independent donors. One-way ANOVA with Tukey’s post-hoc correction for multiple testing, * p<0.05, ** p< 0.01, *** p<0.001.

## DISCUSSION

In this study, we present a novel construct targeting human cDC1s. To the best of our knowledge, we show for the first time the potential of targeting XCR1 on peripheral blood human cDC1s for inducing antigen-specific CD8^+^ T cell responses to a tumor antigen.

Functional analysis of XCL1(CC3) has shown dependency on the integrity of its *N*-, but not C-, terminus to bind XCR1^16^. XCL1 C-terminus can thus be used to introduce a site-specific modification site. Here, a functional chemokine, XCL1(CC3), stabilized in its XCR1 agonist fold^20^, and equipped with a C-terminal LPETGG sortag motif was designed and produced. C-terminal site-specific labeling of chemokines with small probes (< 2 kDa) has been reported^28–30^. Therefore, conjugation of a fluorophore or a tumor antigen might be possible without disrupting its function. XCL1(CC3) selective binding to its target after conjugation to a small fluorophore was confirmed. SDS-PAGE analysis showed that XCL1(CC3) remained partially unreacted, indicating unfolded or misfolded XCL1(CC3), as detected by HPLC. Purity of the final product could be ensured by incubation with anti-FLAG beads, removing unreacted XCL1(CC3).

HLA-A*02:01 is a common haplotype, with a frequency averaging 45% in Europe depending on the ethnicity^31, 32^, and therefore often exploited as model for (cancer) vaccines. The cancer testis antigen NY-ESO-1 contains an HLA-A*02:01 epitope (NY-ESO-1(157-165), ^157^SLLMWITQ^165^C, in short S7C) known to elicit CD8^+^ T cell responses in patients^25^. This peptide has low affinity for HLA-A*02:01 and has a cysteine anchor residue in P9 (^165^Cys). This C-terminal cysteine causes problems in vaccine formulation due to oxidation. This prompted researchers to develop analogs of S7C^26, 33^ in efforts to generate more efficient cancer vaccines. Interestingly, variants of S7C displaying higher affinity for HLA-A*02:01 (^157^SLLMWITQ^165^V, ^157^SLLMWITQ^165^L)^33^ did not show increased immunogenicity, and were not able to elicit cytotoxic T lymphocytes (CTLs) recognizing endogenously processed NY-ESO-1 (157-165). Substitution of the ^165^Cys for ^165^Abu, despite demonstrating weaker binding to HLA-A*02:01, showed increased immunogenicity on patient-derived CD8^+^ T cells ^26^, thus standing out as an ideal vaccine epitope.

While XCL1(CC3)-K(N_3_)-S7Abu was formed in reaction, we were not able to stabilize it in solution, in spite of our efforts (cf. methods, for instance addition of glycerol^34^, or L-arginine HCl solution^35^). S7C and its analogs are notoriously insoluble (solubility around 0.5 mM in DMSO, insoluble in ddH_2_O and Tris-Cl buffer). The peptide was designed with an azido-lysine upstream of the epitope to have the flexibility for further functionalization via SPAAC; which was used to conjugate a 5 kDa PEG chain. PEGylation of proteins is a classical technique to increase hydrophilicity of poorly soluble compounds^36, 37^. Excesses of triazole-PEG5k and K(PEG5k)-S7Abu were removed by cation exchange, and XCL1(CC3)-K(PEG)-S7Abu was purified with over 40% yield, with no detectable aggregation. Importantly, XCL1(CC3)-K(PEG)-S7Abu retained binding to XCR1, likely due to the site-specificity of the conjugation method. Moreover, primary human cDC1s exposed to XCL1(CC3)-K(PEG)-S7Abu were able to elicit antigen-specific CD8^+^ T cell proliferation and activation, testifying that 5 kDa PEGylation did not prevent antigen processing and subsequent presentation on HLA-A*02:01 molecules.

TCR contact residues of immunogenic epitopes tend to be hydrophobic^38^, which is suggested to be a mechanism by which T cells discriminate immunogenic from tolerated epitopes^39^. This is consistent with observations that class I HLA-binding epitopes have a tendency to adopt native α-helical structures^40, 41^, which are amphiphilic with a hydrophobic core. However, short polypeptides typically do not display α-helical structures in solution, as it is not entropically favorable. Therefore, while it is not surprising for short tumor antigens to be poorly soluble, it causes significant challenges in protein-based vaccine formulation. Here, a simple strategy was used, which enhanced peptide solubility and enabled conjugation to a targeting moiety, while not hindering binding, processing and presentation by cDC1s. We hypothesize that this strategy could be extended beyond the model epitope S7Abu.

Importantly, XCL1(CC3)-K(PEG)-S7Abu retained chemotactic activity *in vitro* comparable to rhXCL1 and XCL1(CC3), demonstrating that site-specific modification and PEGylation of XCL1(CC3) did not impair chemokine functionality*. In vivo*, chemokines use GAG binding to establish gradients^43, 44^. The underlying mechanism regulating GAG binding by chemokine oligomers^43^ versus GPCR binding remains unknown. It is hypothesized that either GAG degradation by enzymes, and/or a change in concentration and equilibrium, regulates the local chemokine concentration. XCL1(CC3) in its α-β monomeric fold is reported to have low affinity for heparin, a member of the GAG family^18, 20^. However, it remains very positively charged and could still interact with negatively charged GAGs, and thus generate chemokine gradients *in vivo*. To study whether this is the case, and whether it matters for therapeutic efficacy, a murine model or more elaborate *in vitro* model would be required. We hypothesize that co-administration of soluble FMS-like tyrosine kinase 3 ligand (FLT3L) would attract and induce differentiation of XCR1^+^ DCs^45^, thus increasing the activity of the vaccine.

Data gathered from CD8^+^ T cell activation assays revealed a dose-dependent activity of the constructs. XCL1(CC3)-K(PEG)-S7Abu was already active at low concentrations, and at 0.1 μM, displayed an increased activity compared to sdAb-K(PEG)-S7Abu. At 1 μM, CD8+ T cell activation was enhanced, but both constructs induced a comparable response, as one would expect by incubating phagocytic cells with high amounts of soluble antigens. These results revealed a clear advantage of the XCL1(CC3) construct for targeted vaccination purposes. Notably, our data shows that conjugate uptake is (partially) XCR1-mediated, and that targeting XCR1 on human cDC1s increases the efficacy of antigen presentation. These results are in line with previous research showing that targeting XCR1 *in vivo* increases efficacy of antigen presentation^13, 46^. Additionally, our data supports a previous report showing the efficacy of a hXCL1-glypican DNA-based vaccine^15^, able to chemo-attract lymph node-resident human cDC1s, which in turn activate antigen-specific T cells. DNA-based vaccines are advantageous due to their ease of manufacture; however, their low immunogenicity remains a key bottleneck^47^. Moreover, such a platform allows fusion of XCL1 to a hydrophilic protein, but conjugation to insoluble peptides, such as S7C, is not possible.

In conclusion, we produced a recombinant human XCR1 agonist (XCL1(CC3)), and developed a strategy to conjugate it to a poorly soluble epitope of a clinically relevant tumor antigen. We believe that this strategy is broadly applicable to other insoluble antigens, and will solve a common formulation problem of protein-based fusion vaccines. The fusion construct retained XCR1 binding on primary human cDC1s and chemo-attractive properties. Moreover, cDC1s treated with XCL1(CC3)-K(PEG)-S7Abu induced potent CD8^+^ T cell proliferation and activation. Our results show the potential of XCR1 as a promising target on peripheral blood human cDC1s to induce antigen-specific CD8^+^ T cell immune responses.

## MATERIALS AND METHODS

### XCL1(CC3) design and cloning

pET28a(+) backbone with NdeI and EcoRI cloning sites was obtained by PCR of an existing template (Sec22b in pET28a(+), provided by Dr. Martin ter Beest) using 5’– CCGAGTCACTCATATGGCTGCCGCGCGGCACC –3’ and 5’–ATACATACGAGAATTCGCGGCCGCACTCGAGCACCA –3’. 6xHis-SUMO-XCL1(CC3)-FLAG was ordered as a gBlock from IDT DNA Technologies and cloned in digested pET28a(+) at NdeI (R0111, New England Biolabs) and EcoRI (R0101, New England Biolabs) sites.

### Chemicals

Chemicals used to produce XCL1, sdAb and sortase, used in peptide synthesis and for HPLC purification and analysis, and for cell isolation were obtained from Merck/Sigma Aldrich, unless stated otherwise.

### XCL1 production and purification

XCL1(CC3) was produced in BL21(DE3) *E. coli* by adapting a published protocol^20^. Bacteria were thawed on ice, and incubated for 5 min with 100 ng of XCL1 in pET28a(+) plasmid. Transformation was performed by heat shock in a water bath set at 42°C for 42 sec, followed by 3 min incubation on ice. 1 mL LB was added, and bacteria were left to recover (1 h, 37°C, 220 rpm). Bacteria were transferred to 50 mL of selective media (2xTY + 50 µg.mL^-1^ kanamycin) and grown overnight at 30°C, 220 rpm. The next day, the suspension was inoculated in selective media to OD_600_ ∼ 0.05, and bacteria were grown at 37°C, 220 rpm, until reaching OD_600_ ∼ 0.4–0.8. XCL1 production was induced by addition of Isopropyl β-D-1-thiogalactopyranoside (IPTG) to a concentration of 1 mM, and culturing for 5 h at 30°C, 220 rpm. Bacteria were collected by centrifugation, supernatant was discarded, and pellets were frozen at −20°C until further isolation. Pellets were thawed at room temperature (RT), and resuspended in lysis buffer [50 mM sodium phosphate pH=8.0, 300 mM NaCl, 10 mM imidazole, 20 mM DTT, 20 µg.mL^-1^ of protease inhibitor cocktail (4693159001, Roche)] (10 mL per 1 L original culture). Suspension was sonicated on ice (3 cycles of 30 sec, 25% amplitude), and spun down (8600 g, 30 min, 4°C). Supernatant (soluble fraction) was collected, and pellet was resuspended in resuspension/wash buffer [50 mM sodium phosphate pH=8.0, 6 M guanidine hydrochloride, 300 mM NaCl, 10 mM imidazole, 20 mM DTT] (10 mL per 1 L of original culture) to lyse inclusion bodies. After 1 h of incubation at 37°C on a shaker, suspension was spun down (8600 g, 30 min, 4°C). Supernatant (insoluble fraction) was collected, and pooled with the soluble fraction. Solution was incubated with 2 mL of pre-washed Ni-NTA beads for 1 h at RT. Suspension was diluted 1:1 (*v/v*) with resuspension/wash buffer and transferred to disposable columns. Beads were washed with 25 CV wash buffer, and eluted with 4 CV elution buffer [100 mM sodium acetate pH=4.5, 6 M guanidine HCl, 300 mM NaCl, 10 mM imidazole]. Elution was refolded by infinite dilution in refolding buffer [20 mM Tris pH=8.0, 200 mM NaCl, 10 mM cysteine, 0.5 mM cystine] overnight at RT. Refolded mixture was concentrated on a 10 kDa Amicon spin filter by several cycles of 10 min, 3000 g, RT, until reaching less than 2 mL of volume. SUMO-XCL1 was then diluted with [20 mM Tris pH=8.0], to reach a NaCl concentration of 200 mM, and incubated with ULP-1 SUMO protease (SAE0067-2500UN, Sigma-Aldrich) overnight at RT on an end-over-end shaker. The reaction was subsequently incubated with Ni-NTA beads to remove 6_x_His-SUMO and ULP-1 for 1 h at RT, and transferred to a disposable chromatography column. Flow through was collected, and concentrated on a 3 kDa spin filter at 4°C. Concentration was measured on a Nanodrop 2000 (ND-2000C, ThermoFisher) (MW= 12.1 kDa, ε_280nm_= 8730 M^-1^cm^-1^), and aliquots were stored at −80°C. Production steps, and final product purity were assessed on a 15% sodium-dodecyl-sulfate polyacrylamide gel electrophoresis (SDS-PAGE) using SYPRO Ruby Protein Gel Stain (S12000, Thermo Fisher Scientific). Refolding efficacy was measured by denaturing XCL1 with 20 µM beta-mercaptoethanol (β-ME), and comparing retention times by reverse phase HPLC (C_18_ column).

### Sortase production

Sortase was produced in BL21(DE3) *E. coli* as reported^21^. Briefly, chemically competent BL21(DE3) were transformed by heat shock, and grown overnight at 30 °C in selective media (LB + 50 µg.mL^-1^ kanamycin). The next day, selective media was inoculated at OD_600_ ∼ 0.05, and bacteria were grown at 37°C, 220 rpm to an OD_600_ ∼ 0.6, and induced with IPTG at a final concentration of 1 mM for 16 h at 25 °C. Bacteria were collected by centrifugation, the pellet washed with [50 mM Tris pH=7.5, 150 mM NaCl], and frozen at −20 °C overnight. Pellets were thawed on ice, and resuspended in lysis buffer [50 mM Tris pH=7.5, 150 mM NaCl, 10 mM imidazole, 20 µg.mL^-1^ of protease inhibitor cocktail, 10% (*v/v*) glycerol]. Suspension was lysed by sonication on ice (3×30 sec, 25% amplitude), and centrifuged (8600 g, 30 min, 4 °C). Supernatant was collected, and protein was isolated using Ni-NTA beads. After 1 h of incubation at 4 °C, beads were washed with 100 CV of ice-cold wash buffer [50 mM Tris pH=7.5, 150 mM NaCl, 10 mM imidazole]. Protein was eluted using 2×4 CV ice-cold elution buffer [50 mM Tris pH=7.5, 150 mM NaCl, 500 mM imidazole, 10% (*v/v*) glycerol], and washed by ultracentrifugation at 4°C using a 3 kDa filter (UFC900324, Merck-Millipore) to remove imidazole. Protein concentration was measured on Nanodrop 2000 (MW=17752 Da, ε_280nm_=14565 M^-1^ cm^-1^), and sortase was stored at −80 °C in sortase buffer [50 mM Tris pH=7.5, 150 mM NaCl] supplemented with 10% (*v/v*) glycerol. Protein purity was assessed on a 12% SDS-PAGE. Protein was stored at −80°C.

### sdAb production

A single-domain antibody (sdAb) against BDCA-2 was identified and characterized in house. BL21 (DE3) *E. coli* bacteria were transformed by heat shock and grown overnight at 30°C in 50 mL of LB supplemented with 100 µg.mL^-1^ ampicillin. The next day, bacteria were diluted in fresh selective media, and grown at 37°C to OD_600_∼0.6-0.8. sdAb production was induced with IPTG at a final concentration of 1 mM, and bacteria were cultured overnight at 30°C. Bacteria were pelleted (3000 g, 30 min, 4°C), and resuspended in 20 mL 1×TES [50 mM Tris, 500 mM sucrose, 0.65 mM EDTA] for 1 h at 4°C on a roller shaker. Osmotic shock was performed by diluting suspension with 80mL 0.25×TES and incubated overnight at 4°C. Suspension was spun at 13,000 g at 4°C for 25 min, and supernatant (periplasmic extract) was collected. Periplasmic fraction was incubated with 1 mL pre-washed Ni-NTA agarose (30230, Qiagen) for 1 h at 4°C. Resin was transferred to a disposable column (7321010, Bio-Rad), and washed with 100 CV ice-cold wash buffer [50 mM Tris pH=8.0, 150 mM NaCl, 10 mM imidazole]. Protein was eluted with 2×4 CV ice-cold elution buffer [50 mM Tris pH=8.0, 150 mM NaCl, 500 mM imidazole]. Buffer exchange to sortase buffer [50 mM Tris pH=7.5, 150 mM NaCl] was performed by spin filtration (C7719, Merck-Millipore). Protein concentration was measured using a Nanodrop 2000 (MW= 15.6 kDa, ε_280nm_= 21550 M^-1^cm^-1^)), and purity was assessed by SDS-PAGE. Protein was stored at −80°C.

### Peptide synthesis

[H-GGGCK(FITC)-NH_2_] and [H-GGGK(N_3_)-NH_2_] synthesis was described elsewhere^48^. For [GGGK(N_3_)FRSLLMWITQ(Abu)] synthesis, 25 mL polypropylene syringes with a porous disc were used for solid-phase peptide synthesis (SPPS). In short, 2-chlorotrityl chloride resin (100-200 mesh) was used with a loading of 1.5 mmol.g^-1^ and peptide couplings were performed with a mixture of Fmoc-AA-OH/DIPCDI/Oxyma Pure (3 equiv. each). After cleavage with TFA/TIS/ddH_2_O (95/2.5/2.5), the peptide was precipitated in Et2O, lyophilized and finally purified using a preparative-HPLC-ESI system (Waters) using a C18 reverse phase column, and gradient from 80:20 to 55:45 ddH_2_O:acetonitrile (ACN) in 15 min. Peptide masses were initially predicted by ChemDraw Professional 15.0 (PerkinElmer, Massachusetts, USA) and compared to acquired masses after LC-MS measurements.

### GGG-K(N_3_)-S7Abu conjugation to PEG-DBCO

[GGG-K(N_3_)-FR-SLLMWITQ(Abu)] was dissolved in 1:1 (*v/v*) ACN:ddH_2_O. DBCO-PEG5k (3 equiv.) were added, and reaction was incubated at 37°C for 3 h, and overnight at RT. Conversion rate was assessed by HPLC. Because of cross-reactivity of the thiol in the active site of the sortase with DBCO, free DBCO groups were quenched by addition of 2 molar equivalents of NaN_3_ overnight at RT. Mixture was diluted with ddH_2_O to reduce ACN content to 15% (*v/v*), freeze dried, and lyophilized. Peptide was resuspended at 30 mM in DMSO and used in site-specific modification reactions.

### Analytical HPLC

Analytical HPLC was carried out on Shimadzu instrument composed of a CBM-20A communication Bus module, DGU-20AS degasser, 2 LC20AD pumps, SIL-20AC autosampler, SPD-M20A diode array detector and CTO-20AC column oven. Linear gradient from 5 to 95 % of ACN (+0.036% TFA) into ddH_2_O (+0.045% TFA) were run at flow rate of 1 mL.min^-1^ over 37 min. XSelect® Peptide CSHTM C18, 130Å, 3.5 µm, 4.6 mm × 100 mm (Waters) column was used for the analysis. Chromatographic peak area was determined at λ=220 nm.

### MALDI-TOF

*m/z* of the [GGG-K(N_3_)-FR-SLLMWITQ(Abu)] peptide before and after PEGylation was analyzed on Bruker Microflex LRF MALDI-TOF equipment. Samples were analyzed within a concentration range of 0.3-0.5 mg.mL^-1^ in 1:1 (*v/v*) ACN:ddH_2_O, using 1 µL each of matrix-sample-matrix sown on the MALDI plate. Matrix: Sinapic acid (trans-3,5-dimethoxy-4-hydroxycinnamic acid) (D7927, Sigma Aldrich) at 10 mg.mL^-1^ in ACN:ddH_2_O (1:1 *v/v*) + 0.1 % TFA.

### Troubleshooting NY-ESO conjugation

We aimed at improving the stability of XCL1-K(N3)-S7Abu by modifying several conditions in the workflow (table 1). We notably modified the reaction and purification buffer, which did not change the outcome. We additionally kept the SUMO solubility domain during XCL1 labeling, and aimed at performing ULP1 cleavage afterwards to improve overall solubility during the purification, but the product likewise aggregated before ULP1 digestion.

**Table 1.**
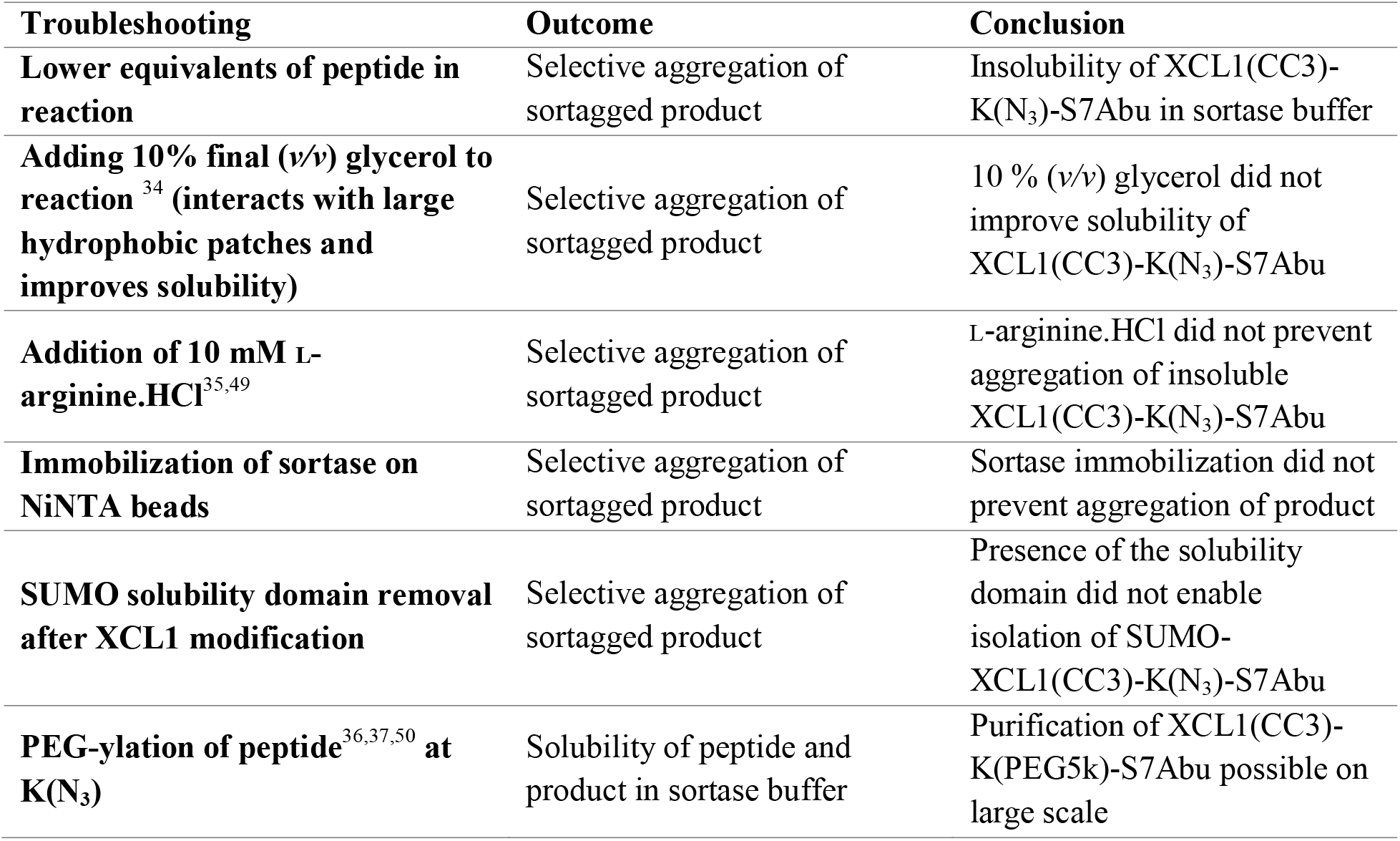
Troubleshooting of site-specific GGG-K(N_3_)-S7Abu conjugation to XCL1.

### Sortagging and purification

Site-specific chemoenzymatic transpeptidation was achieved by incubating XCL1 (final concentration 5 µM) or sdAb (final concentration 20 µM) with 0.75 molar equivalent of 3M sortase A (0.75 equiv.) or 5M sortase A (0.4 equiv.), [H-GGG-X] peptid (25 equiv.), CaCl_2_ (10 mM), DMSO (10% *v/v*), in sortase buffer [50 mM Tris pH=7.5, 150 mM NaCl]. Reactions were incubated at 37°C for 1 h for the 3M sortase, or 2 h at 4°C for the 5M sortase. Reactions were transferred to an appropriate volume of pre-washed Ni-NTA beads and incubated for 30 min at RT on an end-over-end shaker. Reaction was spun over a 100 µm filter to remove Ni-NTA beads, flow-through was collected, and spun down for 10 min at 10 000 g to remove eventual aggregates. Subsequent protein purification workflows are described below. After purification, proteins were concentrated on a 0.5 mL 10 kDa filter (UFC501024, Merck-Millipore) by ultracentrifugation and purity was controlled by SDS-PAGE.

For conjugation to [H-GGGCK(FITC)-NH_2_], product was purified by fast protein liquid chromatography (FPLC, NGC Quest, BioRad) on an ENrichTM SEC70 10×300 column (7801070, BioRad) in sortase buffer.

For conjugation to Cy5.5, XCL1 or sdAb were site-specifically labelled with [H-GGGK(N_3_)-NH_2_], and purified by FPLC to remove peptide excess. XCL1-K(N_3_) and sdAb-K(N_3_) were subsequently reacted with DBCO-Cy5.5 (3 equiv.) (CLK-1046, Jena BioSciences) overnight at RT on an end-over-end shaker. Peptide excess was removed by running the reaction through a PD-10 desalting column (17085101, Cytiva).

For conjugation to [GGGK(PEG5k)FRSLLMWITQ(Abu)], product was loaded onto a strong cation exchange column (HiTrap SP FF, 17505401, Cytiva), and eluted with a gradient of 50 mM Tris + 150 mM – 1 M NaCl. pH of the Tris buffer was adjusted to 7.5 at RT (XCL1 V21C/V59C), or 5.1 at RT (sdAb) to accommodate for the isoelectric point of the protein. Fractions were collected and NaCl concentration was adjusted by diluting with 50 mM Tris to 150 mM. If necessary, protein was incubated overnight at RT with anti-FLAG resin (L00432, Genscript) to remove unreacted XCL1. Protein was concentrated on 3 kDa spin filters, buffer exchange to sortase buffer [50 mM Tris pH=7.5, 150 mM NaCl] was performed, and protein was incubated overnight at 4°C with a Pierce endotoxin removal column (88274, ThermoFisher). Protein was eluted by spin filtration, further concentrated on a 3 kDa filter to reach a final concentration of 1-3 mg.mL^-1^, and aliquoted at −80°C until further use. Endotoxin levels were confirmed below detection levels by chromogenic Limulus amebocyte lysate test, performed by the Radboudumc pharmacy (Nijmegen, NL).

### PBMC isolation from buffycoats

Apheresis material from HLA-A*02:01^+^ donors were obtained from Sanquin (Nijmegen, NL). PBMC suspension was diluted to 300 mL with RT dilution buffer [PBS + 2 mM EDTA] and split over 10×50 mL conical tubes. 10 mL of Lymphoprep (07851, Stemcell) was carefully pipetted underneath, and cells were spun for 20 min at RT, 2100 rpm, brake (2,0). After centrifugation, peripheral blood mononuclear cells (PBMCs) were carefully collected with a 5 mL pipet and transferred to new 50 mL tubes. Cells were diluted up to 50 mL with dilution buffer and spun at 1800 rpm, 10 min, RT. Pellets were washed with 40 mL ice-cold wash buffer [PBS + 1 % human serum albumin (HSA) + 2 mM EDTA] three times, and split over two tubes. Cells were washed one time with ice-cold PBS, and erythrocytes lysed with 10 mL of ACK lysis buffer [150 mM NH_4_Cl, 10 mM KHCO_3_, 0.1 mM Na_2_EDTA] per tube for no more than 5 min at RT. Wash buffer was added to 50 mL, cells were spun down at 1500 rpm, 4°C, 5 min, and resuspended in 50 mL wash buffer. Cells were counted using Türks reagent.

### CD8^+^ T cell isolation and transfection

From each donor, 3–5×10^8^ PBMCs were used to obtain CD8^+^ T cells by negative magnetic isolation (130-096-495, Miltenyi Biotech), according to the manufacturer’s instructions. Cells were counted (expected yield is 5% of starting cells) and washed with a large volume of RT PBS. 10×10^6^ CD8^+^ T cells were resuspended in 250 µL of RT red phenol-free serum-free TheraPEAK^TM^ X-VIVO^TM^-15 (BEBP02-061Q, Lonza). 10 µg of RNA coding for the α and β chains of the TCR recognizing the S7C epitope of NY-ESO-1 presented on HLA-A*02:01 (NY-ESO (157-165)) was thawed on ice, and added to the cell suspension. Cells were transferred to the electroporation cuvette (1652088, BioRad), and transfected using a Gene Pulser Xcell electroporation system (1652661, BioRad) using the following settings: square wave, 500 V, 3 ms, 1 pulse, 4 mm. Cells were carefully transferred to a 15 mL tube containing 1 mL of pre-warmed RPMI + 2 % human serum (HS), and left to recover at 37°C for at least 2 h until further processing. Cells were washed with a large volume of PBS and resuspended in 200 µL PBS + 2% fetal bovine serum (FBS), to which 200 µL PBS + 10 µM Cell Trace Far Red was added. Cells were incubated at 37°C for 20 min, with gentle shaking after 10 min. After 20 min, 2 mL of pre-warmed FBS were added, and cells were further incubated at 37°C for 10 min. Cells were spun down, washed with X-VIVO^TM^-15 (BE02-060Q, Lonza) + 2% HS, and counted (expected yield is 5-6×10^6^ cells). Cells were resuspended in X-VIVO-15 + 2% HS for further use. TCR expression was analyzed the next day by flow cytometry using an anti-mouse TCR antibody (H57-597, BV421, BioLegend).

### Isolation of cDC1s dendritic cells from HLA-A2^+^ apheresis material

From each donor, the rest of PBMCs (typically 2–4×10^9^ cells) was used to proceed to cDC1 isolation. Cell pellet was resuspended in 1 µL per 1×10^6^ cells of microbeads against CD19 (130-050-301, Miltenyi Biotech), CD3 (130-050-101, Miltenyi Biotech), CD14 (130-050-201, Miltenyi Biotech), CD56 (130-050-401, Miltenyi Biotech) and 1 µL per 1×10^6^ cells of FcR block (130-059-901, Miltenyi Biotech). Cell suspension was incubated for 30 min at 4°C on a roller shaker. After 30 min, cells were diluted with a large volume of wash buffer [PBS + 1 % HSA + 2 mM EDTA] and spun down at 1500 rpm, 5 min, 4°C. Cells were resuspended in 24 mL wash buffer, and ran through pre-washed LD columns. Negative fraction was collected and cells were counted. Cells were spun down and resuspended in 1 mL per 1×10^8^ cells, and ran over pre-washed LD columns (130-042-901, Miltenyi, 1 LD per 1×10^8^ cells). Columns were washed with 3×3 mL of wash buffer. Cells were pooled, and spun down at 1500 rpm, 5 min, 4°C. Negative fraction was collected, and columns were washed with 3×2 mL wash buffer. Cells were pooled, and counted. At this point, the yield is expected to be 1-5% PBMCs. Cells were resuspended in 2 µL wash buffer per 1×10^6^ cells, and 1 µL anti-CD1c-biotin (130-119-475, Miltenyi Biotech) per 1×10^6^ cells for 15 min at 4°C, with gentle shaking every 5 min. Cells were washed with a large volume of cold wash buffer, and spun down at 1500 rpm, 5 min, 4°C, and incubated with 2 µL of anti-biotin microbeads for another 15 min. Cells were washed, and resuspended in 1 mL wash buffer, and ran through a pre-washed LS column. Column was washed with 3×3 mL wash buffer, and eluted in a 15 mL tube with 5 mL wash buffer. Positively isolated cDC2s were spun down at 1500 rpm, 5 min, 4°C, and resuspended in 500 µL wash buffer. Cells were run through a pre-washed MS column (130-042-201, Miltenyi), washed with 2×1 mL wash buffer, and eluted with 2 mL wash buffer in a 15 mL tube. Cells were immediately counted, and resuspended in X-VIVO^TM^-15 + 2% HS at 1×10^6^ cells.mL^-1^. Negative fraction (cDC1-containing) was further processed by resuspending in 2 µL wash buffer per 1×10^6^ cells, and 1 µL anti-BDCA-3 microbeads (130-090-512, Miltenyi Biotech) per 1×10^6^ cells for 15 min at 4°C, with gentle shaking every 5 min. Cells were washed with a large volume of wash buffer, and spun down at 1500 rpm, 5 min, 4°C. Cells were resuspended in 1 mL wash buffer, and ran through a pre-washed LS column. Column was washed with 3×3 mL wash buffer, and eluted in a 15 mL tube with 5 mL wash buffer. Positively isolated cDC1s were spun down at 1500 rpm, 5 min, 4°C, and resuspended in 500 µL wash buffer. Cells were run through a pre-washed MS column, washed with 2×1 mL wash buffer, and eluted with 1 mL wash buffer in a 15 mL tube. Cells were immediately counted, and resuspended in X-VIVO^TM^-15 + 2% HS at 1×10^6^ cells.mL^-1^.

### cDC1 / CD8^+^ T cell activation assay

cDC1s from HLA-A*02:01^+^ donors were isolated. In a round bottom 96-well plate (3799, Corning), 10 000 cDC1s were seeded in 50 µL, and treated with various conditions for 3 h, in a final volume of 100 µL X-VIVO^TM^-15 + 2% HS + 1 µg.mL^-1^ poly(I:C) (tlrl-picw, Invivogen). After 3 h, 100 µL X-VIVO^TM^-15 + 2% HS was added, and cells were spun down. Supernatant was carefully pipetted off, and cells were resuspended in 100 µL X-VIVO^TM^-15 + 2% HS + 0.6 µg.mL^-1^ poly(I:C). 50 000 TCR-transfected CD8^+^ T cells were added in 100 µL, and cells were incubated for 120 h. 50 µL supernatant was harvested after 24h. Division index was calculated by the formula 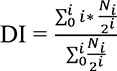, where, i is the division cycle number, and N_i_ the percentage of live CD8^+^ T cells in that cell cycle.

### *In vitro* chemotaxis assay

50 000 freshly isolated primary cDC1s were placed in 100 µL X-Vivo^TM^-15 + 2% HS in the top compartment of a 5µm HTS transwell (3388, Corning). Lower compartment was filled with 200 µL X-Vivo^TM^-15 + 2% HS, eventually supplemented with 10 ng.mL^-1^ recombinant XCL1 (758002, BioLegend), 10 ng.mL^-1^ in house-generated XCL1(CC3), or 10 ng.mL^-1^ XCL1(CC3)-K(PEG)-S7Abu, and cells were incubated for 15 h. Cells migrating towards the lower compartment were directly harvested and counted using a MACSQuant (Miltenyi). XCL1-mediated chemotaxis was calculated by the formula:

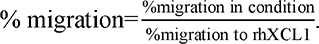

### Flow cytometry

Stainings were performed in 50 µL in 96-well plates. Live/death staining was performed with fixable eFluor506 viability dye (1:2000, 65-0866-14, ThermoFisher) for 30 min at RT in PBS. Antibody stainings were performed in PBA [PBS + 5% FBS + 0.01% NaN_3_] for 30 min at 4°C. DC purity was assessed by antibodies against CD141 (1:10, APC, AD5-14H12, 130-113-314, Miltenyi) and XCR1 (1:10, PE, S15046E, 372604, BioLegend), and T cell phenotype was assessed by antibodies against CD25 (1:100, AF488, BC96, 302616, BioLegend), CD127 (1:100, PE/Cy7, A019D5, 351320, BioLegend), CD137 (1:100, PE, 4B4-1, 555956, BD Pharmigen), CD279 (1:100, BV421, EH12.2H7, 329920, BioLegend), and TIGIT (1:100, BB700, 741182, 747846, BioLegend). Cells were washed 2 times with PBA before acquisition on a FACSLyric (BD).

### ELISA

Enzyme-linked immunosorbent assay (ELISA) was performed for IFNγ and IL-2 using uncoated kits (respectively 88-7316-88 and 88-7025-88, ThermoFisher) following manufacturer’s instructions. Briefly, plates were coated with 100 µL capture antibody diluted in coating buffer [50 mM Na_2_CO_3_/NaHCO_3_, pH=9.6] overnight at 4°C. Plates were washed 3× with wash buffer [PBS + 0.05% Tween-20], blocked for 1 h at RT with 200 µL ELISA diluent reagent, and washed 1×. Supernatants were diluted in ELISA diluent reagent (1:50 for IFNγ and 1:10 for IL-2), and 100 µL were incubated for 2 h at RT with the capture antibody. Plates were washed 5× and incubated with 100 µL diluted detection antibody for 1 h at RT. Plates were washed 5×, and incubated with 100 µL diluted streptavidin-HRP for 30 min at RT. Plates were washed 7×, and incubated with 100 µL TMB. Reactions were developed in the dark, and stopped with 100 µL 2 M H_2_SO_4_. Absorbance was measured at λ = 450 nm on an iMark microplate reader (1681130, Bio-Rad). Standard was prepared in duplicate and used to perform quantification using a Four parameter logistic (4PL) modeling (available at: https://www.arigobio.com/elisa-analysis).

### Microscopy

Freshly isolated cDC1s were resuspended in PBA and stained with either XCL1(CC3)-Cy5.5 or sdAb-Cy5.5 for 30 min at 4°C. Cells were washed 1x with 100 µL PBA and 1x with 100 µL PBS. Cells were resuspended in 100 µL 4% paraformaldehyde (PFA), and fixed for 30 min at 4°C. In the meantime, 12 mm circular confocal slides (72230-01, 1 ½, Electron Microscopy Sciences) were placed on parafilm, cleaned with 100% ethanol for 1 min, and washed 2x with 100 µL PBS. Slides were coated with ±100 µL of 10 µg.mL^-1^ poly-L-lysine in ddH_2_O for 25 min at RT. Cells were spun at 2000 rpm for 10 min at RT, and washed 2x with 100 µL of RT PBS. Slides were washed with PBS and 40 µL of cell suspension was mounted for 30 min at RT in the dark. Finally, slides were mounted face down on 4 µL embedding medium [24% (w/v) glycerol, 9.6% (w/v) Mowiol 4-88, 1.5 μg.mL^-1^ DAPI, 0.1 M Tris pH=8.5], left to polymerize in the dark at RT overnight, and further stored at −20°C. Fluorescence was acquired on a LSM900 (Zeiss, Jena, DE).

## AUTHORS CONTRIBUTIONS

CML, GFG and MV conceived the project and designed experiments with the help of IJMdV and CGF. CML, AC, LdH and GFG performed experiments. CML and AC analyzed the data under supervision from GFG and MV. IRT, JCE, AMDB, ZW and YD contributed experimentally. CML and MV wrote the manuscript with assistance from all authors. All authors reviewed the manuscript.

## ACKNOWLEDGEMENTS

We thank Prof. Dr. Ton N.M. Schumacher (NKI, Amsterdam, NL) for permission to use the TCR sequence, and Prof. Dr. Uğur Şahin and Dr. Mustafa Diken (BioNTech, Mainz, DE) for providing mRNA encoding the TCR recognizing the NY-ESO-1 (157-165) epitope. We thank Dr. Marc Dalod for advice during project design.

## DATA AND MATERIALS AVAILABILITY

All data needed to evaluate the conclusions are present in the paper and/or the Supplementary Materials. The described plasmids used in this study are deposited in plasmid repository of Addgene (www.addgene.org/), or may be requested from the authors

## APPENDIX

**Table 2.**
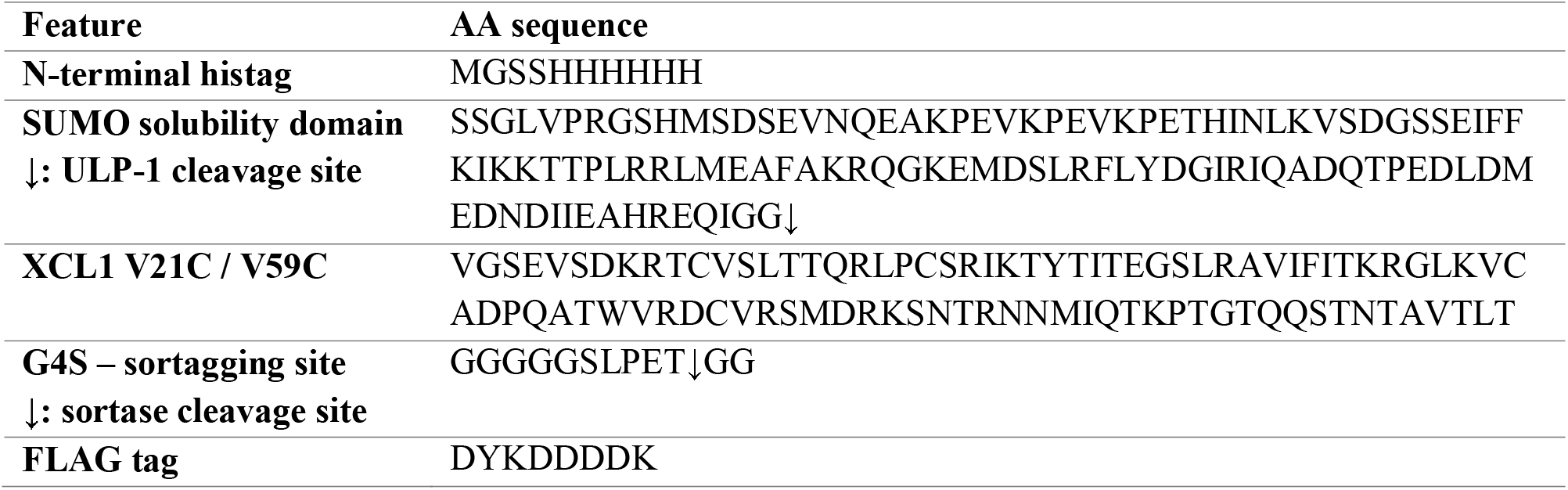
Amino acid sequence of 6×His-SUMO-XCL1-LPETGG-FLAG.

## LIST OF SUPPLEMENTARY MATERIAL

**Figure S1:**
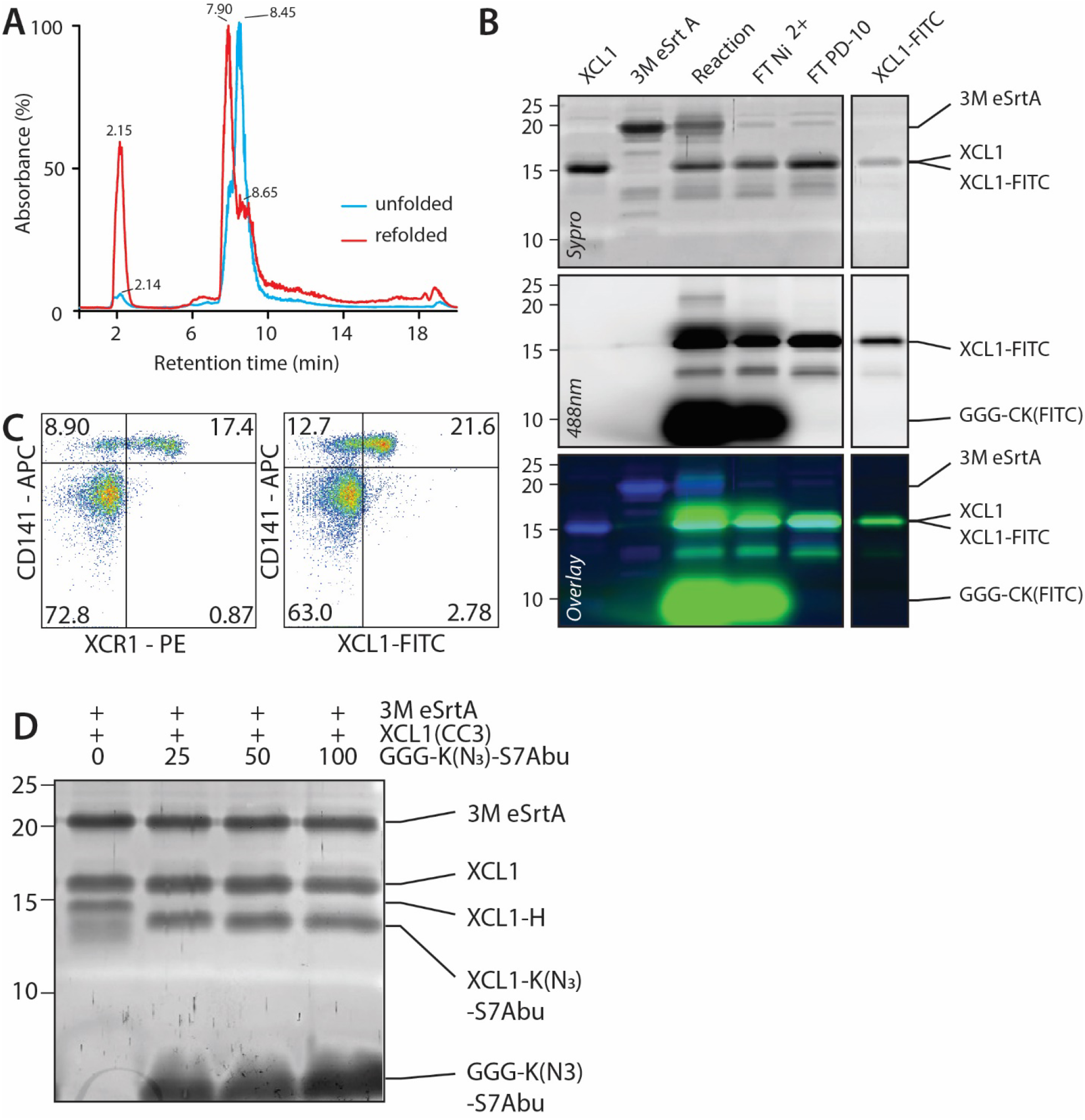
Production, characterization and site-specific labeling of XCL1(CC3) with GGG-K(N_3_)-S7Abu. **A.** HPLC analysis of XCL1(CC3) (refolded, red) and β-ME-treated XCL1(CC3) (unfolded, blue) shows a distinct retention time of XCL1(CC3) upon refolding. Refolding rate was calculated by comparing the ratio of the two species to be over 75%. **B.** Site-specific labeling of XCL1(CC3) (lane 1) with 3M eSrtA (lane 2). After reaction (lane 3), XCL1(CC3)-FITC is purified by nickel affinity purification (FT Ni^2+^, lane 4), FITC excess removed by PD-10 desalting (lane 5) and pure product is obtained after concentration (lane 6). **C.** Commercial anti-XCR1-PE and XCL1(CC3)-FITC identify a comparable subpopulation of ∼60% CD141^+^ cDC1s without staining CD141^-^ cDC2s, confirming that XCL1(CC3)-FITC specifically binds to XCR1. **D.** Small-scale optimization of GGG-K(N_3_)-S7Abu peptide equivalents for site-specific labeling of XCL1(CC3) shows that only 25 equivalents of GGG-K(N_3_)-S7Abu are necessary to generate XCL1(CC3)-K(N_3_)-S7Abu with minimal hydrolysis byproduct (XCL1-H).

**Figure S2:**
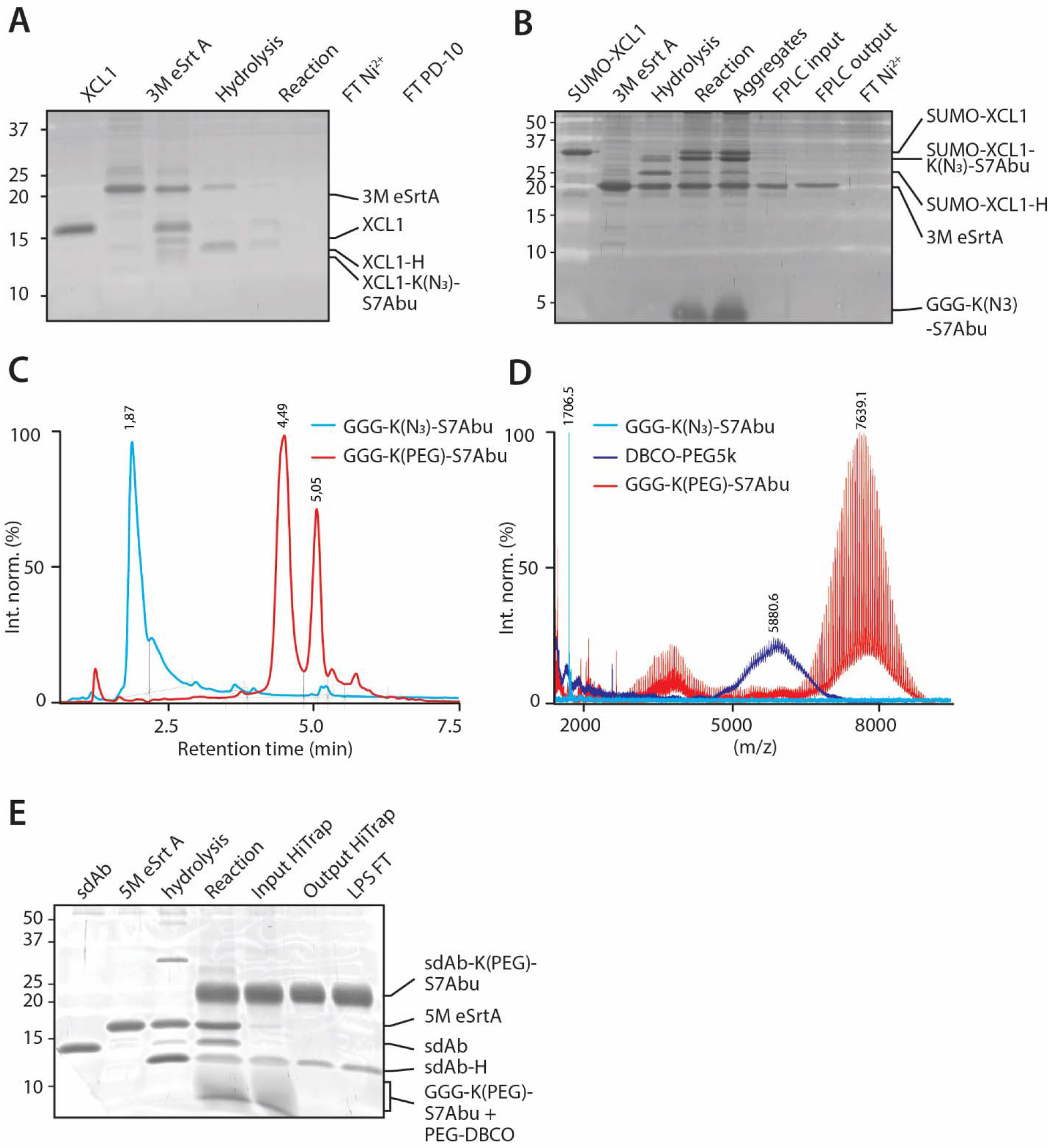
PEGylation of K(N_3_)-S7Abu enables purification of stable XCL1(CC3)-K(PEG)-S7Abu. **A.** Site-specific labeling of XCL1(CC3) (lane 1) with GGG-K(N_3_)-S7Abu allows XCL1(CC3)-K(N_3_)-S7Abu product formation (lane 4), but product could not be purified (lanes 5 and 6). **B.** Site-specific labeling of SUMO-XCL1 (XCL1 fused to its solubility domain) with GGG-K(N_3_)-S7Abu allows product formation (lane 4), but product is present in aggregates formed during reaction (lane 5), and could not be isolated (lanes 6-8). **C.** HPLC analysis of GGG-K(N_3_)-S7Abu before and after PEGylation shows a distinct retention time, and reaction completion. **D.** MALDI-TOF analysis of GGG-K(N_3_)-S7Abu, DBCO-PEG5k and GGG-K(PEG)-S7Abu shows an increase in *(m/z)* upon PEGylation, and reaction completion. **E.** sdAb-K(PEG)-S7Abu can be generated and purified by cation exchange, with minimal hydrolysis product present. Densitometry was performed to calculate the concentration of sdAb-K(PEG)-S7Abu used in cell experiments.

**Figure S3:**
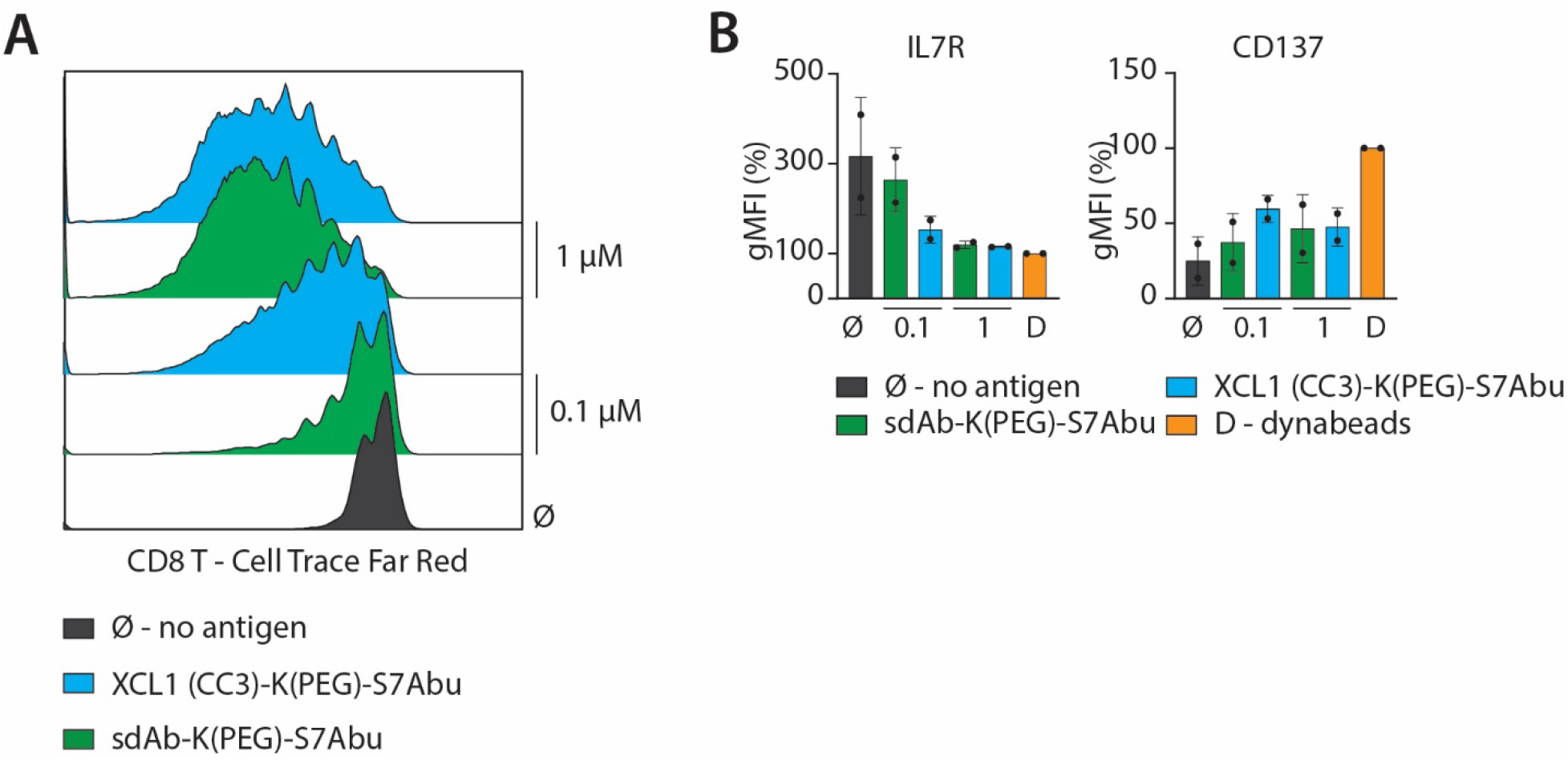
XCL1(CC3)-K(PEG)-S7Abu is degraded by cDC1s, and S7Abu presented to CD8^+^ T cells. **A.** CD8^+^ T cell proliferation tracked by cell trace dye upon treatment of cDC1s with XCL1(CC3)-K(PEG)-S7Abu or sdAb-K(PEG)-S7Abu shows the selective advantage of XCR1 targeting. Representative of N=3 independent donors. **B.** IL7R and CD137 expression on activated CD8^+^ T cells. N=2 independent donors.

